# Effect of rectified gap junctional electrical coupling and spatial distribution of biologically engineered pacemaking cells on ventricular excitation

**DOI:** 10.1101/2022.12.02.518808

**Authors:** Yacong Li, Qince Li, Jun Liu, Lei Ma, Kuanquan Wang, Henggui Zhang

**Affiliations:** School of Computer Science and Technology, Harbin Institute of Technology, Harbin, China; Beijing Academy of Artificial Intelligence, Beijing, China; Peng Cheng Laboratory, Shenzhen, China; National Biomedical Imaging Center, Peking University, Beijing, China; Biological Physics Group, School of Physics and Astronomy, The University of Manchester, Manchester, United Kingdom

**Author notes:** Correspondence: Qince Li: Room 810, Zonghe Building, No. 92, West Dazhi Street, Harbin, China, Henggui Zhang: Room 3.07, Schuster Building, Brunswick Street, Manchester M13 9PL.

**Keywords:** Biological pacemaker, Gap junction, Rectifying property, Cardiac modelling

## Abstract

**Aim:** Biologically engineered pacemaker, or bio-pacemaker, is a promising replacement for electronic pacemakers for treating cardiac dysfunction. Previous animal experimental studies, however, have not been able to accurately demonstrate the stability and efficiency of the bio-pacemaker yet. This study aimed to elucidate the underlying factors that affect bio-pacemaker’s performance and to discover possible optimising solutions to enable the potential use of bio-pacemaker therapy.

**Methods and results:** The human ventricular myocytes model in this study followed the ten Tussucher’s model in 2006, and the bio-pacemaker single cell model was modified based on it as what has been expatiated in our previous work. In tissue model, two factors were primarily evaluated for their effects on bio-pacemakers to pace and drive surrounding cardiac tissue: gap junction between bio-pacemaker cells (PMs) and adjacent ventricular myocytes (VMs) and the spatial distribution of bio-pacemakers. A suppressed gap junctional electrical coupling between and heterotypic gap junctions were simulated and a combination of them led to the best performance of the bio-pacemaker. Then, the pacemaking behaviours of three kinds of idealised PM-VM slices were simulated, in which an electrically isolated distribution of bio-pacemaker showed optimal drive capacities. Finally, a real human ventricular slice model was used to verified the conclusions in idealized tissues.

**Conclusion:** This study develops a theory that weak-rectified electrical coupling and electrically isolated distribution can enhance the pacemaking efficiency of bio-pacemakers, which lays the groundwork for future research into therapeutic applications of bio-pacemakers.

**Author summary:** Biologically engineered pacemakers are expected to be a substitute for electronic pacemakers because of their physiological superiority, but how to transform them for practical application remains challenging. In this paper, we presented a theoretical perspective on optimising biological pacemaking capability based on a computational simulation approach. By manipulating the gap junctional electrical coupling among bio-pacemaking cells and between the pacemaker and their surrounding cells, and controlling spatial distribution of bio-pacemaker, we demonstrated that an enhanced capacity of a bio-pacemaker can be achieved. The results of this study may provide a theoretical basis for the further clinical development of bio-pacemakers.

## 1. Introduction

The biologically engineered pacemaker (bio-pacemaker) has been presented as a new potential for the treatment of sick sinus syndrome and atrioventricular block (1-3). Superior to the electronic pacemaker, the bio-pacemaker has been believed to be sensitive to emotion and hormones (4, 5) and thus may provide a better prognosis for patients. There was some evidence implying that non-rhythmic cardiac myocytes can be transformed into bio-pacemaking cells (PMs) by gene engineering (6-8), and their underlying pacemaking mechanisms were analysed in some simulation studies (9-11). Although the stability of single pacemaker cells still needs to be optimised in biological experiments, these studies provide single cell foundations for the bio-pacemaker therapy.

Animal studies have shown some potential in this area. Plotnikov et al. (12) injected HCN2-induced human mesenchymal stem cells (hMSCs) into the canine left ventricular anterior wall to drive the heart in complete heart block. An *in vivo* experiment found that injecting the Tbx18 adenoviral vector into the apex of a guinea pig’s heart can produce ectopic pacemaker activity when sinus rhythm was suppressed by methacholine (6). A similar method also worked on the large animal. Tbx18 viral vectors can maintain the heart rate of a pig with complete heart block. And the Tbx18-induced porcine heart was sensitive to autonomic regulation and did not present pro-arrhythmic risk in the short term (13).

Previous biological experiments demonstrated the possibility of the clinical use of a bio-pacemaker, but there remain some key questions that must be further elucidated. Particularly, it should be understood which factors have an influence on the drive capacity (i.e., how many cardiac cells that the bio-pacemaker can drive) of the bio-pacemaker and how they impact the generation and propagation of spontaneous pacemaking signals. In the sinoatrial node (SAN), the native cardiac pacemaking cell, two factors help to maintain the source-sink balance: a weak cell-cell gap junction between SAN cells and insulating spatial construction by the atrium (14). These features of the natural cardiac pacemaker can guide the design of artificial bio-pacemakers.

A normal SAN tissue has two functions: (1) ensuring inherent automaticity; (2) bearing the hyperpolarised voltage load of adjacent atrial cells and conducting the pulse to the atrium (15). The particular connexin of gap junctions in the SAN helped to maintain its functions (16). In the SAN, connexin 45 (Cx45) is mainly expressed while atrial myocytes (AMs) primarily express connexin 43 (Cx43) (17, 18). The conductance of Cx45 is less than Cx43 (19), which means that the gap junctional electrical coupling in the SAN is far below that of normal cardiac myocytes (CMs). In this way, the inherent automaticity of SAN can be preserved from the depression by adjacent cells. In bio-pacemakers, a two-cell unit experiment (20) showed that Cx43 was expressed to establish the gap junction between PMs and CMs so that the pacemaking signal could propagate. Meanwhile, the downregulation of Cx43 was observed in Tbx18-induced pacemaker cells (13). Hence, a threshold of connexin expression must exist in PMs to balance the generation and propagation of spontaneous signals.

The specific coupling approach between the SAN and the atrium is as crucial as the gap junction in the SAN tissue to ensure the pacemaking function of SAN. For SAN, the atrium is equal to an electrical load (21, 22). However, anatomically, approximately 10,000 SAN cells (23) can drive the whole heart with billions of non-rhythmic cells (24). There are three hypotheses of the combination mode between SAN and the atrium: mosaic theory, gradient theory, and rectification theory. The mosaic theory states that SAN and atrium are randomly distributed at the junction (25). Gradient theory claims that the size and electrical properties of SAN cells change gradually from centre to periphery (26, 27). In rectification theory, there is a diode-like structure of electrical coupling between SAN and adjacent CMs via co-expressed Cx45/Cx43 (28, 29), i.e., heterotypic gap junction, which suppresses the current from AMs to SAN while promoting the reverse current (15). According to our previous simulation research, the gradient model describes the characteristic electrical activity of SAN better than the mosaic model (30), and the gradient SAN-AM model has been used to investigate the effect of fibroblasts (31) or SCN5A mutations (32) on wave conduction in the heart. The rectified property appeared in heterotypic gap junctions which were composed of two different hemichannels (33), while general homotypic gap junction consists of two identical hemichannels. In the heart, gap junctions are usually encoded by Cx40, Cx43 and Cx45 (34). HeLa cell experiments verified that these three kinds of connexin can be compatible with a functional gap junction channel and present rectified conductance (19, 35). According to the possible SAN characteristics (28, 29), we hypothesised that PMs and CMs can be coupled in a rectified way, by which the hyperpolarised potential of non-rhythmic CMs was hindered in bio-pacemakers so that their inherent pacemaking ability could be played to the best.

Except for gap junctions, the special coupling approach between SAN and AMs is also reflected in the specific spatial distribution of SAN and the atrium. The SAN is separated from the atria by connective tissue (27, 36), which serves as a natural barrier to block the hyperpolarised potential of AMs. Similar functional role of conduction barrier was also observed in an engineered bio-pacemaker experiment. A biological experiment (5) revealed that reducing the cell-cell coupling and physical connection of a bio-pacemaker with neighbouring cardiac tissue could ensure spontaneous pacemaking waves to propagate continually. According to this experiment, an isolated distribution would be applied to construct a bio-pacemaker model and be compared with other distributions.

The location of the bio-pacemaker in the heart may also affect its pacemaking capacity. The electronic pacemaker is implanted in the right atrium and the right ventricle depending on the different indications, sick sinus syndrome and atrioventricular block, respectively (37). To avoid desynchrony of the ventricular electro-mechanical coupling function, electrodes can be implanted in the ventricular septum or the Hickey tract (38). In animal experiments, bio-pacemakers have also been located in the right ventricle to drive the heart (6, 13), but the underlying impact of bio-pacemaker location on the synchronising duration of tissue needs further research.

In an electrically isolated bio-pacemaker experiment, a pacemaker cell could only drive 13 CMs in average (5), with a pacemaking capacity far below that of SAN. This number was only two in a one-dimensional cardiac tissue model with reduced boundary coupling between the PMs and adjacent ventricular myocytes (VMs) (39). Obviously, the drive ability of current engineered bio-pacemakers still needs to be improved.

Thus far, several key factors that may affect the bio-pacemaker’s performance have been outlined, but they have yet to be quantitatively linked and analysed. Our study aims to determine the effects of cell-cell electrical coupling and spatial distribution on the drive capacity of a bio-pacemaker by constructing a cardiac tissue model consisting of a bio-pacemaker and ventricular tissue. In this paper, heterotypic gap junctions with rectified properties were modelled and applied between PMs and VMs in a cardiac model. Combining the rectified electrical coupling (REC) model between PMs and VMs with a weak electrical coupling (WEC) model among PMs, the pacemaker performance was enhanced dramatically. In an isolated pacemaker implantation schedule with WEC and REC based on a real ventricular structure, one pacemaker could drive more than 800 VMs on average, which indicated that it was possible to improve the drive capacity of artificial bio-pacemakers by modifying their gap junction and spatial distribution. This paper aims to pave the way for the experimental explorations of bio-pacemakers.

## 2. Methods

### 2.1 WEC model

According to the nonlinear cable theory, the isotropic two-dimensional cardiac tissue model can be expressed by the following partial differential equation (40):

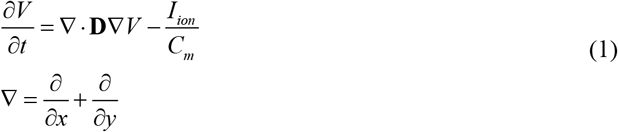

where *V* is the membrane potential; *t* is time; **D** is a tensor of the diffusion coefficient; *C*_*m*_ is cell capacitance; *I*_*ion*_ is the sum of transmembrane ionic currents.

The *I*_*ion*_ of PMs and VMs can be referenced in (9) and (41), respectively. Particularly, in PM model, we set the conductance of the inward rectifier potassium channel current (*G*_*K1*_) at 0.54 nS/pF and the conductance of the funny current (*G*_*fNa*_, *G*_*fK*_) at (0.047 nS/pF, 0.0279 nS/pF), so that the pacemaking activity of single PM cells was stable at a cycle length of 853 ms.

An idealised spatial distribution of the cardiac tissue model is shown in Figure 1, which consisted of PMs and VMs. The heterogeneous ventricular tissue was fixed at 3 × 100 cells, including endocardial cells, middle cells and epicardial cells (ENDO:MCELL:EPI) with a ratio of 25:35:40.

**Figure 1.**
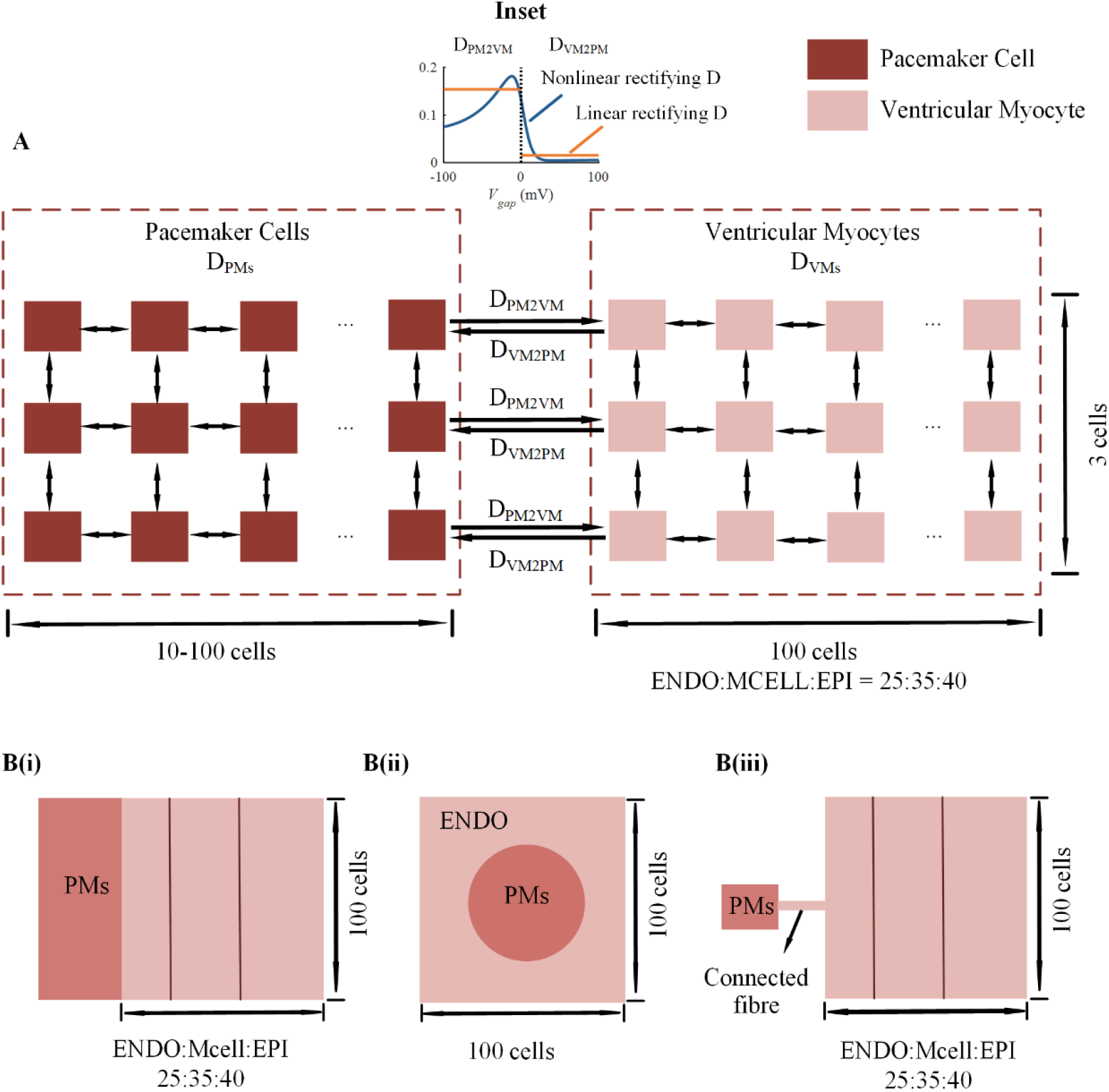
Spacial distribution of cardiac tissue model and electrical coupling structure. A: Illustration of diffusion coefficient (**D**) in a full side-to-side coupling model. B: Spatial distribution of cells in a full side-to-side coupling model, an embedded model and an isolated model. Inset: The value of D between PM and VM.

In the cardiac tissue model, **D** determines the coupling level of the gap junction between cells and affects the propagation of the electric wave in tissue. In the present model, the diffusion coefficient was heterogeneous. We defined the value of **D** between PMs as D_PMs_, the value of **D** between VMs as D_VMs_, the value of **D** from PM to VM as D_PM2VM_ and the value of **D** from VM to PM as D_VM2PM_, which are illustrated in Figure 1A.

In the WEC model, D_VMs_ was set at 0.154 mm^2^/ms (the same as that in original ventricular tissue (41)), as well as D_PM2VM_ and D_VM2PM_, whereas D_PMs_ was reduced to simulate the downregulation of gap junctions in PMs. The length of PMs was set as a variable to evaluate the drive capacity of the bio-pacemaker under different D_PMs_ values.

### 2.2 REC model

#### 2.2.1 Linear REC model

In basal cardiac tissue model, **D** between two cells was a fixed value. Here, we set **D** between PM and VM as a piecewise function to simulate the rectified property between cells (dashed line, Figure 1 Inset):

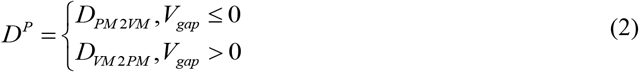

in which *D*^*P*^ is the value of the diffusion coefficient between PM and VM; *V*_*gap*_ is defined as the voltage difference between PM and VM as follows:

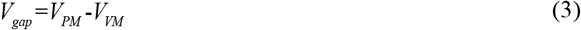

In this model, the gap junctional current between PM and VM was asymmetric with respect to *V*_*gap*_ = 0. When D_VM2PM_ was less than D_PM2VM_, the conductance from VM to PM was suppressed, whereas the opposite conductance was promoted. In this way, the effect of the VMs’ hyperpolarising potential on PM was suppressed, so the intrinsic membrane potential of PMs was preserved. When D_VM2PM_ was greater than D_PM2VM_, the conductance of gap current from VM to PM was greater than the reverse. When D_PM2VM_ is equal to D_VM2PM_, the model was returned to the original.

#### 2.2.2 Nonlinear REC model

The simplified linear rectified model was suitable for revealing the effect of REC on the production and propagation of automatic pacemaking signals. However, biological experiments demonstrated that the conductance of heterotypic gap junctions was a nonlinear function of the voltage difference between two cells (*V*_*gap*_) (19). According to a HeLa cell experiment (42), the relationship between rectified electrical coupling and the voltage difference of two consecutive cells can be fitted by a Boltzmann equation. Here, we built a rectified diffusion coefficient model as follows:

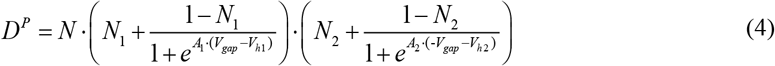

in which *N, N*_1_, *A*_1_, *V*_*h*1_, *N*_2_, *V*_2_, and *V*_*h*2_ are Boltzmann constants; *V*_*gap*_ is defined in Eq. (3).

Boltzmann constants were set as Eq. (5) to fit the rectified electrical coupling shape in Ref. (42) with a maximum D value close to that of ventricular tissue. This situation was defined as original rectified *D*^*P*^ as shown in the inset of Figure 1.

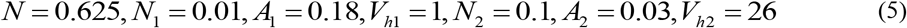

Substituting Eq. (4) into Eq.(1) makes the two-dimensional cardiac tissue model be further solved as follows:

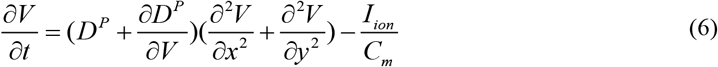

The solving process and numerical solution were presented in S1 Text.

### 2.3 Spatial distributions of bio-pacemaker in cardiac tissue model

Three idealised spatial distributions were designed in this study. A full side-to-side coupling model was designed in which a block of rectangular pacemaker tissue clung to a piece of heterogeneous ventricular tissue (Figure 1B(i)). The triple-layer of ventricular tissue was set as ENDO:MCELL:EPI = 25:35:40. In the embedded model, a round pacemaker tissue was embedded in a square cardiac tissue at 100 × 100 cells (Figure 1B(ii)). In the isolated model, pacemaker tissue was connected with 100 × 100 triple-layer heterogeneous ventricular tissue through ENDO fibres as shown in Figure 1B(iii). Apart from the three idealised models, a real human ventricle structure derived from American virtual body data was used to simulate the implantation scheme of a bio-pacemaker in the heart. The left and right ventricles were included in this slice with a total size of 436 × 461 cells.

In each spatial structure, the electric activities of cardiac tissue under normal and weak-rectified electrical coupling (W-REC) conditions were simulated to study the effect of electrical coupling and spatial distribution on the drive capacity. The drive capacity is defined as follows:

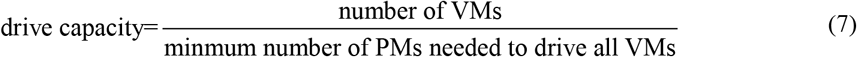

Drive capacity represents the average number of ventricular cells that can be driven by a single pacemaker cell. It can be influenced by many factors, such as the inherent automaticity of PMs, the electrical coupling mode of PMs with surrounding cells, and the spatial distribution of pacemakers in cardiac tissue. Therefore, the value of drive capacity itself can only reflect the driving ability of a specific cardiac tissue, which was used to estimate the pacemaking ability under control. The higher the drive capacity was, the stronger the pacemaker’s drive ability.

## 3. Results

### 3.1 Effect of WEC between PMs on bio-pacemaker performance

WEC was simulated by reducing D_PMs_. Figure 2A shows the effects of downregulating connexin expression in PMs (i.e., D_PMs_) on the initiation and conduction of pacemaker activity in the model with the structure defined in Figure 1A. In the figure, the percentage of D_PMs_ was used for presentation with 100% D_PMs_ equal to the value in human ventricular tissue model (41). The average number of PMs that can drive 100 VMs (i.e., the length of PM tissue in Figure 1A) was used to evaluate the drive capacity of the bio-pacemaker. The lower the number of PMs was used, the better the drive capacity was. Figure 2A shows that with 100% D_PMs_, at least 37 PMs were needed to drive 100 VMs. When the D_PMs_ was suppressed to 20%, the minimum number of PMs that could activate 100 VMs decreased to 19. The results suggested that PMs number required to drive VMs declined almost linearly with the reduction in D_PMs_. The present simulation results quantifiably verified the positive effect of decreasing connexin between PMs on the bio-pacemaker’s performance, but it was still far away from the drive capacity of SAN. Therefore, the WEC bio-pacemaker model should be further optimised to satisfy physiological needs.

**Figure 2.**
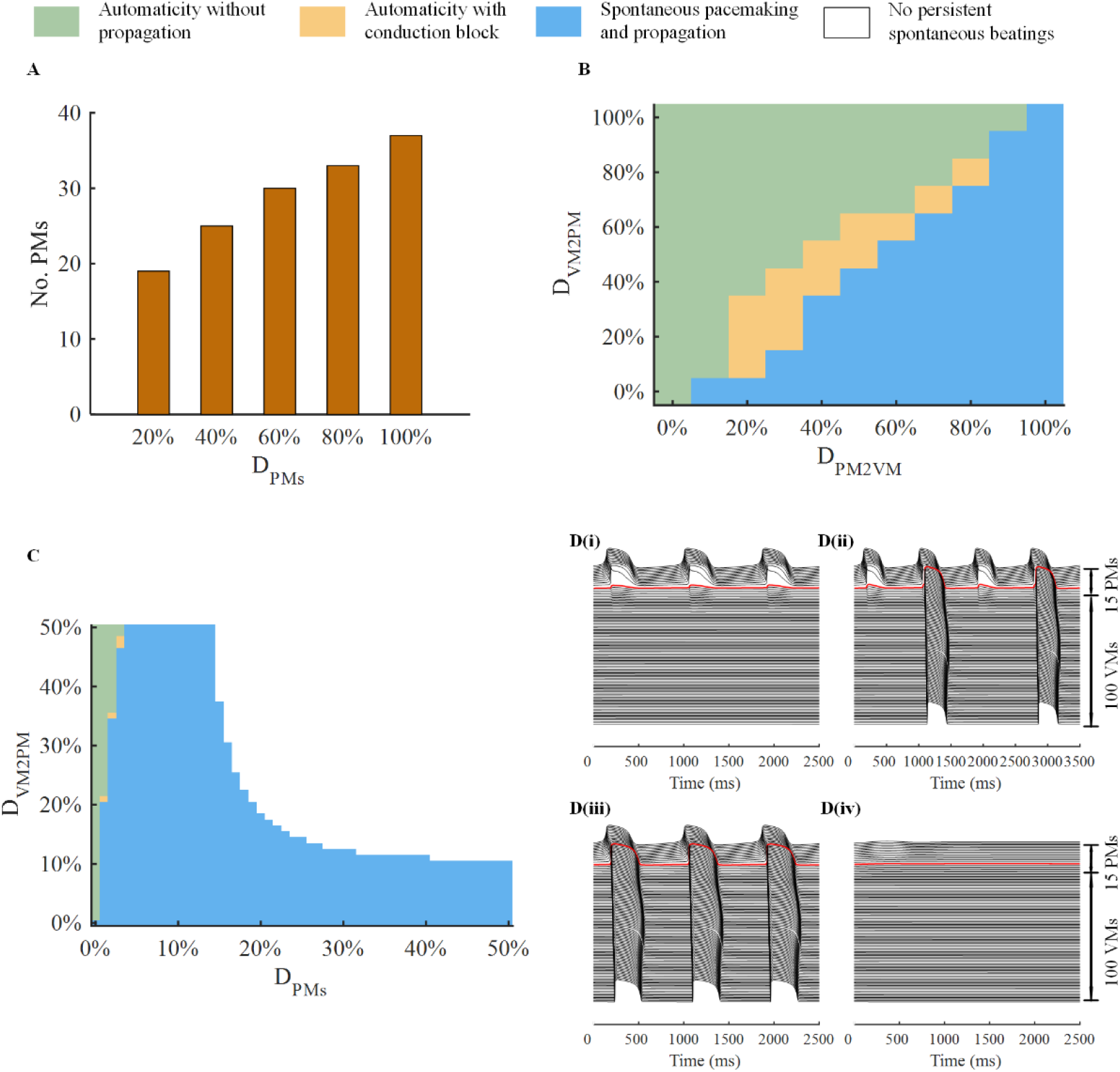
Effect of W-REC on pacemaking states in idealised cardiac tissue. A: Average minimum numbers needed to drive 100 VM cells under different D_PMs_. B: Pacemaking states of cardiac tissue under different D_PM2VM_ and D_VM2PM_ percentages. C: States of electrical excitation waves in cardiac tissue under different D_PMs_ and D_VM2PM_ percentages. White: No persistent spontaneous beatings; Green: Automaticity without propagation; Yellow: Automaticity with conduction block; Blue: Spontaneous pacemaking and propagation. D: Sequence diagrams of the membrane potential of a row of horizontal continuous cardiac cells when (D_PMs_, D_VM2PM_) = (20%, 30%), (2%, 40%), (2%, 35%), (2%, 20%). The red line marks the membrane potential of the PM neighbouring VM tissue.

### 3.2 Effect of linear REC on bio-pacemaker performance

Linear REC model was conducted by modifying D_PM2VM_ and D_VM2PM_ in Eq. (2), by which the rectified direction and degree were varied. The tissue size was fixed at 15 × 3 PM cells and 100 × 3 VM cells. The evaluation index for drive capacity was whether PM tissue can produce spontaneous electrical waves that propagated into adjacent VM tissue.

When cell-cell coupling between PMs was kept at its original value (D_PMs_ was 100%), the bio-pacemaker can initiate spontaneous waves that propagate into VM tissue only when D_PM2VM_ and D_VM2PM_ were 100% and 0% respectively (figure not given).

Furthermore, the rectified property was studied under the condition of WEC. Figure 2B illustrates the electrical signal states of tissue under variable D_PM2VM_ and D_VM2PM_ when D_PMs_ was reduced to 10%. There were three kinds of states: (1) automaticity that cannot be conducted: spontaneous activation of PMs occurred but could not propagate into VM tissue (marked by green in Figure 2B); (2) automaticity with conduction block: the automaticity of PMs could partially propagate into VM tissue (marked by yellow in Figure 2B); and (3) conductible automaticity: the automaticity of PMs could totally propagate into VMs (marked by blue in Figure 2B). When D_VM2PM_ was equal to or greater than D_PM2VM_ (green and yellow area), it was difficult for the pacemaking signals to propagate into the VMs. When D_VM2PM_ was less than D_PM2VM_ (blue area), depolarising signals were induced spontaneously and propagated into adjacent ventricular tissue. Additionally, as the rectified level (the difference between D_VM2PM_ and D_PM2VM_) increased, the pacemaking cycle length (PCL) shortened (Supplementary Figure S1A). Simulation results tentatively verified the positive effect of the rectified property on the pacemaker.

### 3.3 Combined effect of WEC and REC on bio-pacemaker performance

Simulations results shown above suggest that REC can improve pacemaking performance in combination with WEC conditions. We further investigated the integrated effects of WEC and REC on the generation and conduction of spontaneous pacemaking waves.

The simulations were conducted by using the spatial distribution of cardiac tissue as shown in Figure 1A, in which the PM tissue size was fixed at 15 × 3 cells. Three parameters were considered in simulations: the D_PMs_ that reflected the WEC level between PMs, D_VM2PM_ that reflected REC level between PMs and VMs and D_PM2VM_ which was fixed at an original value to maintain the electrical conductivity from PMs to VMs. Simulation results showed that the electrical exciting waves can be classified into four states in the whole parameter space of D_PMs_ and D_VM2PM_ as shown in Figure 2C, and each representative state of action potentials in a row of horizontal continuous cells is presented in Figure 2D. In Figure 2D(i), although PMs could produce automaticity, these waves could not spread into adjacent VM tissue, which is marked as green in Figure 2C. In the yellow parameter space of Figure 2C, the electrical exciting waves can propagate into VM tissue but are sometimes blocked, as shown in Figure 2D(ii). In Figure 2D(iii), automatic pacemaking signals can propagate into VM tissue completely and the parameters are marked as blue in Figure 2C. In the white space of Figure 2C, there was no depolarising behaviour in cardiac tissue as shown in Figure 2D(iv).

Figure 2C reveals that the states of electrical exciting waves were related to the degree of D_PMs_ and D_VM2PM_. When D_PMs_ = 0, there was no electrical coupling between PMs, so spontaneous beating could not propagate into the ventricle (Figure 2D(i)). When D_VM2PM_ was large (between 15%-50%), a small D_PMs_ could not generate pacemaking activity in tissue because the cell-cell coupling between PMs was too weak to transmit depolarising potentials into VM tissue (green area in Figure 2C and Figure 2D(i)). A slight increase in D_PMs_ or decrease in D_VM2PM_ could partially propagate spontaneous electrical signals into the ventricular tissue (yellow area in Figure 2C and Figure 2D(ii)). Less than 20% of D_PMs_ or less than 15% of D_VM2PM_ could ensure that every spontaneous pulse in PMs propagated into VM tissue (blue area in Figure 2C and Figure 2D(iii)), and greater D_PMs_ made tissue stay at resting potential throughout the whole simulation period (white in Figure 2C and Figure 2D(iv)). Either small D_PMs_ (less than 20%) or small D_VM2PM_ (less than 15%) extended the other parameter space that could produce synchronous pacemaking behaviour, and the same tendency appeared in a wide parameter space of 0-100% D_PMs_ and D_VM2PM_. When D_VM2PM_ =100%, the scenario was equivalent to the WEC model built in Section 3.1.

Above results indicated that the effects of WEC and REC on the pacemaker–ventricular tissue model were different. The decrease in D_VM2PM_ (i.e., REC condition) was conducive to the conduction of pacemaking behaviour, and the smaller the value of D_VM2PM_, the more likely it was to produce a stable pacemaking state (Figure 2C). Therefore, REC had a monotonic effect on the pacemaking state. However, the situation became different in WEC condition. Reducing D_PMs_ moderately can guarantee pacing behaviour, but insufficient D_PMs_ resulted in conduction block.

The reason why spontaneous beating failed to propagate into adjacent cells at the threshold of incomplete and complete conduction states (yellow and blue areas in Figure 2C) was analysed. Simulation results indicated that a tiny difference in the adjacent current caused by acute electrical coupling resulted in the suppression of depolarisation. Figure 2D shows that the conduction of electric exciting waves was closely related to the potential difference across the border between PM and VM tissue (Figure 2D, red line). In cases where the peripheral PMs were successfully depolarised, the stimulation current offered by PM was sufficient to produce action potentials in ventricular tissue. To explore the subcellular reasons for the automaticity conduction, three cases were analysed: (D_PMs_, D_VM2PM_) = (2%, 17%), (2%, 18%) and (2%, 19%). Figure 3 shows the time sequence diagrams of the action potential, *I*_*adj*_ and *I*_*out*_ of a peripheral PM cell in the different cases. *I*_*adj*_ is defined as the sum of currents from neighbouring cells to the present PM, including depolarising current from PMs (inward current, *I*_*in*_) and repolarising current from VMs (outward current, *I*_*out*_). When D_VM2PM_ was decreased to 19%, *I*_*out*_ was greater because of the strong repolarising effect of VM on PM (Figure 3C). During the repolarisation phase of the first pacemaking behaviour (at approximately 1100 ms), the greater *I*_*out*_ caused a larger *I*_*adj*_, which made the maximum diastolic potential more negative. Before the depolarisation phase of the second pacing cycle (1500–1800 ms), a greater D_VM2PM_ led to a larger load current, which exceeded inward current (|*I*_*out*_|>|*I*_*in*_|). Consequently, *I*_*adj*_ was excessive (Figure 3B), and membrane potential failed to depolarise (Figure 3A).

**Figure 3.**
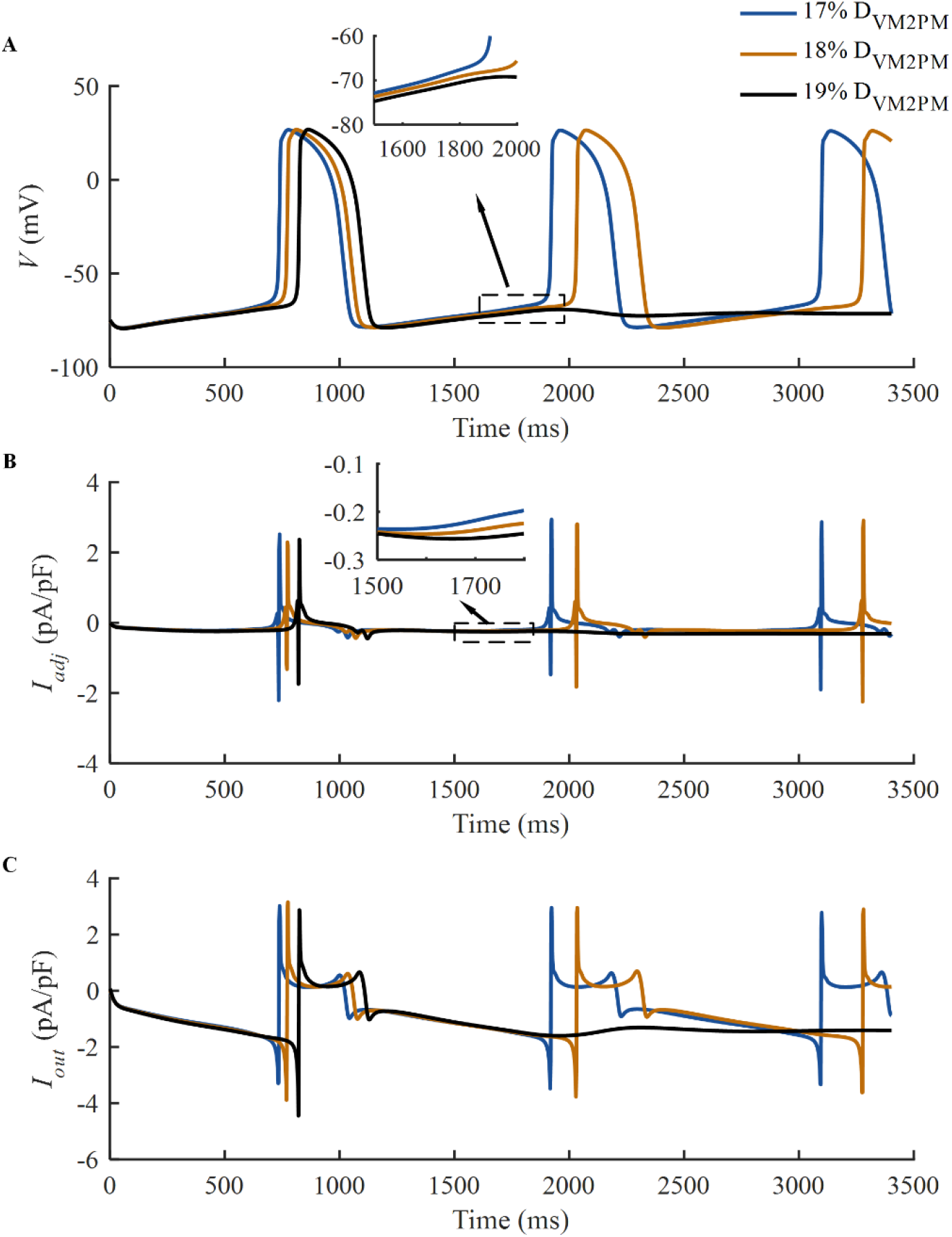
Mechanism of automatic electrical signal propagation from PM to VM. (A-C) *V, I*_*adj*_ and *I*_*out*_ of the PM neighbouring VM tissue when (D_PM2VM_, D_VM2PM_) = (2%, 17%), (2%, 18%), (2%, 19%) respectively.

In addition to pacemaking states, the effects of REC and WEC on electrical conduction properties were also examined by calculating PCL and conduction velocity (CV). PCL was defined as the PCL of the first PM cell and CV was measured across the tissue horizontally from the first VM to the last VM. Statistically, as the D_PMs_ or D_VM2PM_ decreased, the PCL became shorter (Supplementary Figure S1B), demonstrating a nearly monotonic change. CV was mainly affected by D_PMs_ rather than D_VM2PM_. Larger D_PMs_ contributed to a faster CV (Supplementary Figure S1C) and weakened the impact of D_VM2PM_ in CV.

### 3.4 Nonlinear REC model

The *V*_*gap*_–*D*^*P*^ relationship in original nonlinear rectified D model with parameters of Eq. (5) is shown in Figure 4A (blue line). The simulation results showed that there was no spontaneous pacemaking activity that could be provoked in cardiac tissue (Figure 4B(i)). According to the analysis in the linear rectified model, the silence of PM tissue may be caused by the heavy load from VMs. In other words, decreasing *D* when *V*_*gap*_ was greater than 0 probably helped to maintain the automaticity of PMs. Therefore, *A*_*1*_ and *V*_*h1*_ in Eq.(4) were modified to 0.5 and -5, respectively, so that the *V*_*gap*_–*D*^*P*^ curve changed to the orange line in Figure 4A. In this case, automaticity appeared and could propagate into VM tissue (Figure 4 B(ii)).

**Figure 4.**
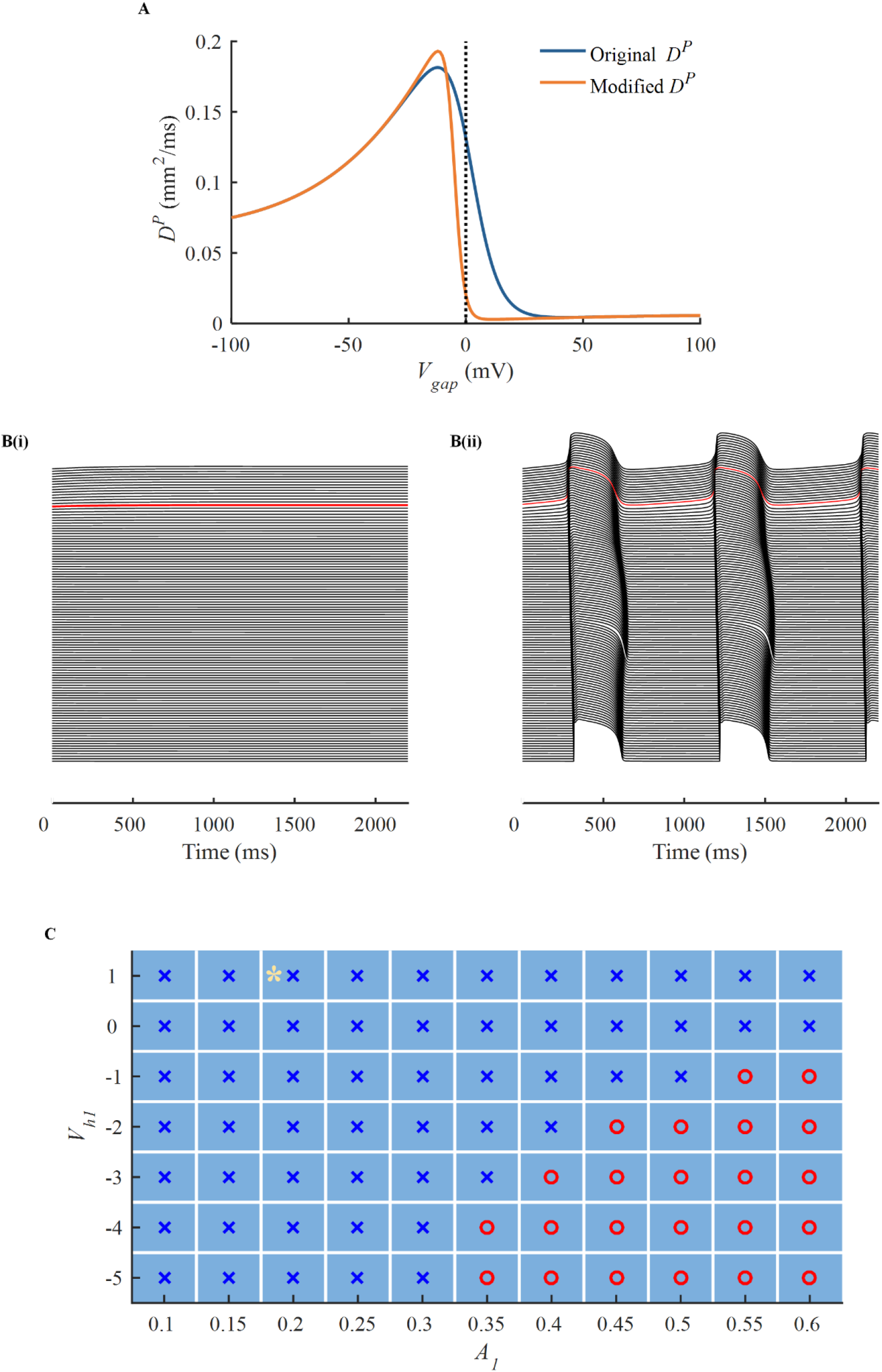
Effect of nonlinear REC on the bio-pacemaker performance. (A) The relationship between *V*_*gap*_ and the diffusion coefficient between PM and VM (*D*^*P*^) in original and modified value with (*A*_*1*_,*V*_*h1*_) is (0.18, 1) and (0.5, -5) respectively. (B) Sequence diagrams of the action potential from a row of horizontal continuous cardiac cells under original and modified nonlinear *D*^*P*^ as shown in Figure A. (C) Pacemaking states of cardiac tissue with different nonlinear REC parameters. The blue cross marks the cases of no normal electrical signal in tissue. The red circle indicates that the pacemaking wave can be propagated into VMs. The yellow asterisk marks the original parameters.

To estimate the effect of nonlinear rectified properties on the pacemaking state of cardiac tissue, we simulated the electrical activities under different *A*_*1*_ *and V*_*h1*_ values. The results are presented in Figure 4C where the original value is marked by a yellow asterisk. The blue cross and red circle mark the cases of abnormal and normal electrical conduction states, respectively. Figure 4C shows that the increase in *A*_*1*_ as well as the decrease in *V*_*h1*_ were necessary to induce spontaneous beatings in cardiac tissue.

We also simulated the bio-pacemaker with nonlinear REC under WEC conditions. When the D_PMs_ was greater than 20%, the parameter space of *A1* and *Vh1* that can produce normal pacemaking behaviour was similar to that under the condition of original D_PMs_ (figure not shown). When D_PMs_ was reduced to 10%, the bio-pacemaker with original nonlinear rectified D model provoked normal pacemaking signals. Hence, the decrease in D_PMs_ can also maintain the ability of bio-pacemakers to pace under the condition of nonlinear rectified D, which was in accordance with the result in Section 3.3.

### 3.5 Effect of spatial distribution on the drive capacity of bio-pacemaker

Three representative types of bio-pacemaker spatial distributions (full side-to-side coupling model, embedded model, and isolated model) were considered to study possible effects of various connection types between PM and VMs on the pacemaking activity in cardiac tissue. The minimum number of PMs that were needed to drive the whole ventricular slice were calculated to assess the pacemaking performance. Both normal cells coupling and W-REC conditions were simulated in each of the distribution models. For the W-REC condition, a linear REC was used when D_PMs_ and D_VM2PM_ were decreased to 5% and 10%, respectively. Depolarisation time was defined as the time from spontaneous signal initiation to whole tissue depolarisation to reflect the depolarisation efficient of cardiac tissue.

In the full side-to-side coupling model, under normal coupling conditions, an average of 37 PMs were needed to drive 100 VMs, whereas only nine PMs were needed in the W-REC model (Figure 5B(i) and C(i)). Representative snapshots from spontaneous pacemaking activity initiation, propagation and repolarisation to the next depolarisation are shown in Supplementary Figure S2. And the depolarisation time was 160 vs. 90 ms in the normal and W-REC models, respectively. Additionally, the PCLs of the two cases were 1280 and 1034 ms, respectively.

**Figure 5.**
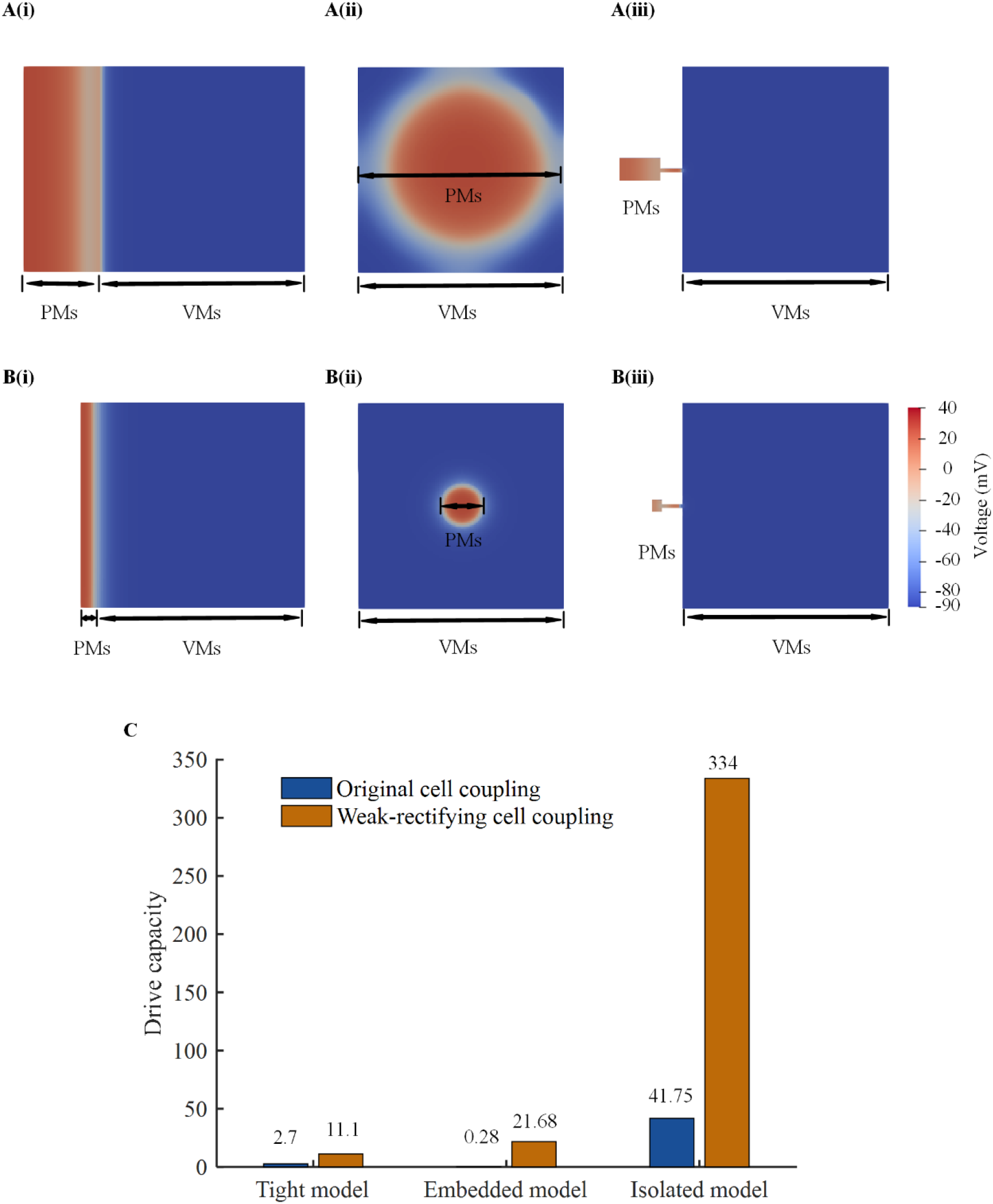
Effect of bio-pacemaker’s spatial distribution on its drive capacity. (A) Initial potential mappings of cardiac tissue under normal electrical coupling conditions with different spatial distributions. (B) Initial potential mappings of cardiac tissue under W-REC conditions with different spatial distributions. (C) The drive capacity of bio-pacemakers under various spatial distributions and electrical coupling modes.

For the embedded distribution model, in the state of normal coupling, a radius of 50 PMs was needed to induce automaticity in cardiac tissue (Figure 5B(ii)), meaning that at least 7843 PMs were needed to drive only 2157 VMs. The W-REC state could decrease the minimal radius to only 12 PMs (Figure 5C(ii)). The process of automatic signal initiation and conduction can be seen in Supplementary Figure S3. The depolarisation time of cardiac tissue on normal and W-REC was 100 and 64 ms, respectively, and the PCL was 1110 and 1090 ms, respectively.

The isolated model with normal coupling needed at least 20 × 12 PMs to drive 100 × 100 VM tissue (Figure 5B(iii)), whereas this number was reduced to only 5 × 6 in the W-REC model (Figure 5C(iii)). The conduction process of spontaneous pacemaking activity is shown in Supplementary Figure S4. The depolarisation time in normal and weak-rectified conditions was 54 and 50 ms, respectively, and the PCL was 1154 and 1092 ms, respectively.

The drive capacities defined in Eq. (7) under different distributions and electrical coupling are compared in Figure 5C. The isolated distribution facilitated pacemaking behaviour significantly. Under the condition of W-REC, the isolated model improved the drive capacity by approximately 30 times when compared with the full side-to-side coupling model and in normal cell coupling condition, a 15-fold improvement appeared. All three types of pacemaker distributions can benefit from W-REC. Particularly, in embedded models, the drive capability increased by 77 times when electrical coupling was modified.

### 3.6 Simulation of bio-pacemaker in a real ventricular tissue model

The present bio-pacemaker was embedded into the right ventricle according to the operation of a porcine experiment (13). The previous results demonstrated that W-REC significantly enhanced pacemaking efficiency. Therefore, the bio-pacemaker was modelled in W-REC mode with the D_PMs_ and D_VM2PM_ reduced to 5% and 10% respectively.

A 15 × 6 isolated bio-pacemaker was inserted into the right ventricle through an isolated distribution, which can initiate spontaneous electrical waves in cardiac tissue. One periodic pacemaking activity of cardiac tissue can be seen in Figure 6A. When the simulation time was 700 ms, PM tissue began depolarising. Then, the electrical wave propagated into adjacent ventricular tissue during 700–868 ms. Following repolarisation, the tissue was placed into a resting state before reactivating at 1580 ms. In this condition, the depolarisation time was 168 ms. The spontaneous electrical waves of the ventricular tissue implanted with isolated bio-pacemaker were shown in Video S1. With side-to-side coupling, at least 10 × 20 PMs can drive the whole ventricular tissue as shown in Supplementary Figure S5.

**Figure 6.**
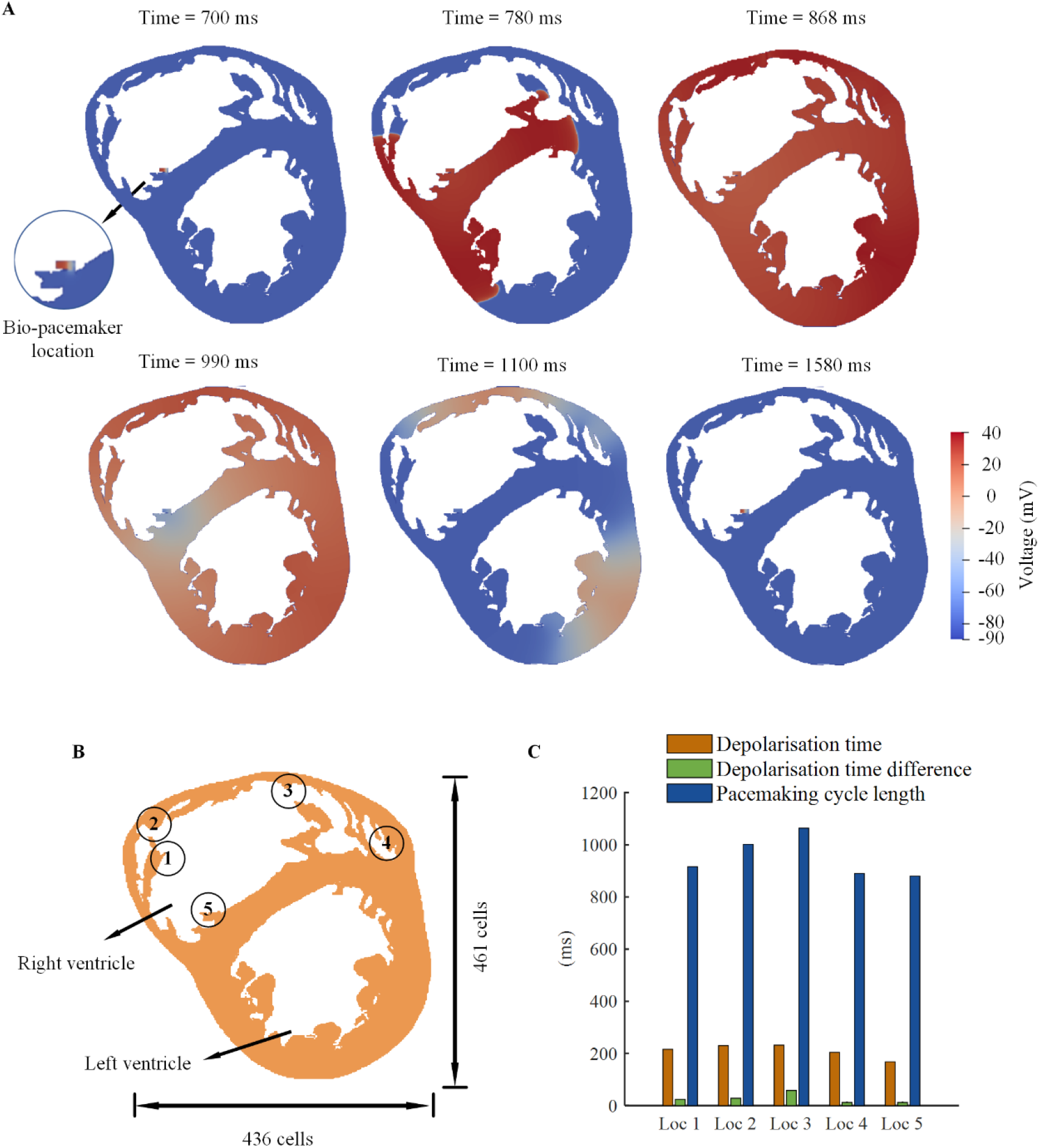
Effect of bio-pacemaker location on the synchrony of cardiac tissue. (A) Potential mappings of cardiac tissue implanted with isolated bio-pacemaker under W-REC conditions. (B) Location of bio-pacemaker in a real cardiac model. (C) Pacemaking properties at different pacemaker locations.

If the left and right ventricles depolarise simultaneously reflects the synchronous activation behaviour. It was defined as the difference between the simulation time that the whole left ventricle reaches depolarisation and that time in right ventricle, which was called depolarisation time difference. The effect of the bio-pacemaker’s location on synchrony was estimated by simulating the pacemaking behaviour of a real ventricle implanted with bio-pacemaker in five different locations (Figure 6B). Based on the simulations, the depolarisation time of tissue, depolarisation time difference, and PCL of cardiac tissue were calculated and are shown in Figure 6C. The results indicated that the bio-pacemaker’s location mainly had effects on depolarisation time and depolarisation time difference rather than PCL. The bio-pacemaker had the shortest depolarisation time and depolarisation time difference at the Loc. 5, i.e., the interventricular area. As a result, it can be inferred that the interventricular area may be an optimal position to place bio-pacemakers so that the left and right ventricles could pace synchronously.

## 4. Discussion

The bio-pacemaker has shown potential as a new therapy for cardiac dysfunction. Related research has mainly focused on the induction of the pacemaking function at the single cell level and has made great progress (43, 44). Animal experiments have created short-term bio-pacemakers in both rodents (6) and mammals (13). However, until now, the conception of the idealised bio-pacemaker in 2004 (3) has not been accomplished, especially as far as its stability and efficiency. In our previous study, a stable single pacemaker cell model was built and the underlying mechanisms that informed the pacemaker robustness were analysed (9). In this study, we systematically investigated the factors that affect the drive efficiency of the bio-pacemaker for the first time, promoting the progress of research and clinical application of bio-pacemaker.

Gap junctions are essential to maintain and propagate the heart rhythm. Abnormal connexin expression may cause heart disease (45), but modulating its expression provides the possibility of optimising the drive ability of bio-pacemakers. Based on the results of experiments showing reduced Cx43 expression in Tbx18-induced pacemaker cells (5, 6, 13), we investigated the relationship between WEC in PMs and drive ability. We then attempted to incorporate heterotypic gap junctions that can be observed biologically (46) into bio-pacemaker therapy design. A heterotypic gap junction has a rectified property, which is expected to maintain the inherent pacemaking ability of PMs as well as propagate spontaneous signals into neighbouring non-rhythmic cells. Our study verified that incorporating WRC and REC can significantly enhance pacemaking efficiency. Moreover, spatial structure was considered in this study because of its importance in native SAN function. In agreement with both SAN (47) and Tbx18-induced pacemaker (5) experimental results, isolated structures preserved the pacemaking rhythm of the bio-pacemaker extremely well in this study. Finally, a real ventricular tissue model study manifested that a lathy shape of bio-pacemaker in ventricular septum was the optimal scheme for the synchronisation of cardiac pacing. In summary, this study provides two major contributions: (1) a novel rectified cell-cell electrical coupling approach for the design of bio-pacemaker is proposed to improve its drive capacity; (2) in real heart tissue model, the spatial distribution of bio-pacemakers and the coupling strategy between bio-pacemakers and VMs are provided to produce the optimal driving capacity.

### 4.1 Gap junction in bio-pacemakers

Prior studies have noted the importance of WEC in bio-pacemakers. VMs can be turned into pacemaker cells, generating spontaneous signals by infecting with Tbx18 via adenoviral vectors (6, 13), not only because of the decrease in I_K1_ and the expression of I_f_ in single PMs but also because of the downregulation of connexin. Therefore, we incorporated WEC in the present bio-pacemaker model.

Rectifying property refers to the unilateral conductivity of a semiconductor. Several reports have shown that rectified property of gap junction can also be observed *in vivo* (48). This type of gap junction provides a voltage-dependent gate with rectified properties, which allows the gap current to flow normally in one direction while being diminished in the other. This is exactly what a bio-pacemaker needs to preserve its own membrane potential from the voltage of neighbouring cells while reserving the ability to transmit its rhythm to them. However, the heterotypic gap junction between bio-pacemaker cells and ventricles was not considered in previous engineered bio-pacemaker studies. As Cx43 and Cx45 are widely expressed in the atrium and SAN, respectively (33), we hypothesised that a heterotypic Cx43–Cx45 gap junction can be constructed between VM and PM. Based on this hypothesis and a HeLa cell experimental data (42), we built a mathematical model to describe a REC between PMs and VMs and simulated its application for bio-pacemaker therapy.

One interesting finding is that the degree of connexin expression that regulates W-REC was crucial to ensuring the normal function of a bio-pacemaker, in terms of both the pacemaking ability and the driving capacity. There was a difference in the effect of WEC and REC on electrical signal propagation in bio-pacemaker model. REC had a monotonic effect on the pacemaking state while WEC’s effect was nonlinear (Figure 2). Increasing the level of REC or WEC can reduce the interference of surrounding cells with PM cells and ensure the inherent pacemaking ability of PM cells, thus decreasing PCL (Supplementary Figure S1). However, low WEC level inhibited the automaticity of PMs transmitting to VMs in space, thus blocking the propagation of electrical waves. In addition, the REC defined in this study mainly affected PMs rather than VMs and had little effect on CV in the VM tissue.

The above simulations were carried out based on the hypothesis that the coupling of cardiac cells was linear. Furthermore, according to the asymmetric property of Cx43–Cx45 gap junctions (42), a nonlinear REC model was developed to better describe the heterotypic gap junctions as shown in Figure 4. According to the biological property of Cx43–Cx45 gap junctions, the modifications of *A*_*1*_ and *V*_*h1*_ are corresponding to the Cx45 hemichannels, i.e., the hemichannels expressed in PMs. Consequently, the present study speculates that re-editing the connexin gene of PMs can optimise bio-pacemaker without changing the genes of adjacent cells.

In conclusion, based on the heterotypic gap junction observed in the HeLa cells, the present study proposed a theoretical possibility to improve bio-pacemaker therapy by designing a heterotypic gap junction. If the technology is fulfilled by gene editing, it may significantly improve the performance of bio-pacemaker and promote the clinical application of bio-pacemaker therapy.

### 4.2 Isolated distribution of bio-pacemaker in cardiac tissue

The physical connection between pacemaker cells and non-rhythmic cardiac cells has been extensively researched in SAN (27, 49) but has rarely been investigated in bio-pacemakers. A bio-engineered research study demonstrated that electrical insulation by neighbouring CMs helped source-sink (spontaneous pacemakers - non-rhythmic cardiac cells) match (5). In this study, using a computational approach, the effect of pacemaker’s spatial distribution on the pacemaking activity were investigated. Simulation results in Figure 5 was consistent with experimental observations (5) that isolated distributions produced the best driving performance in three types of spatial distributions. In the isolated model with normal electrical coupling, the maximum number of VMs that can be paced by a single pacemaker cell was approximately 3 times greater than that in experiment (5). More importantly, under the W-REC condition, the maximum number exceeded 30 times that observed in biological experiment. Hence, re-editing the bio-pacemaker electrical coupling connexin may significantly improve the drive capacity, under both isolated distribution and other distributions.

An animal study has reported that more than 700,000 hMSCs can restore the heartbeat in complete heart block condition, yet fewer hMSCs (257,000) could also work in some circumstances (12). The underlying mechanism of this phenomenon remains mysterious. According to the simulations in this study (Figure 5 and 6), various structure of PMs may account for this phenomenon. Fewer hMSCs may located in the heart by an insulating structure, thus facilitating the generation and preservation of its own automaticity. Though this hypothesis needs further validation via more animal studies, it may provide a novel technique to design bio-pacemaker therapy via changing the spatial structure.

In addition to the best drive capacity, the bio-pacemaker model of isolated distribution also produced the shortest depolarisation time in three kinds of distributions. The conduction velocity was quicker in ventricle than other kinds of cardiac tissues. As the original ventricular structure was preserved in the isolated pacemaker distribution by implanting less extra PM cells, the electrical conduction efficiency of the original ventricular tissue could be maintained. These results demonstrate the potential advantage of W-REC under isolated distribution.

### 4.3 Bio-pacemaker implantation scheme in a real ventricular tissue model

Based on studies in idealised pacemaker-ventricular tissue, we implanted bio-pacemakers in a real ventricular structure using a W-REC mode. The 10 × 20 side-to-side bio-pacemaker as shown in Figure S5 can activate whole ventricular tissue with a PCL of 876 ms when the intrinsic PCL of single PMs was 853 ms (9), which is consistent with the normal rhythm of human VMs. Increasing the size of the pacemaker to 15 × 20 can shorten the PCL to 860 ms (result is not shown). Therefore, if merely increasing the number of pacemakers, a 50% increase in pacemakers is required to decrease the PCL by 1.8%, which was highly inefficient. Consequently, this study suggested increasing pacemaking efficiency by modifying the coupling mode between PMs and VMs rather than by increasing the pacemaker size.

Intriguingly, pacemakers with the same size but different contact areas between PMs and VMs produced different pacemaking states. When the size of bio-pacemaker was 20 × 10 cells (i.e., the contact area between the pacemaker and ventricle has 20 cells), there was no spontaneous pacemaking activity during the whole simulation period (result is not shown). However, the bio-pacemaker with a size of 10 × 20 was able to produce spontaneous pacemaking activity as shown in Figure S5. This suggested that the contact area of the bio-pacemaker played a prominent role in the pacemaking function, which was consistent with the conclusion that electrically isolated spatial distributions could preserve the inherent performance of the bio-pacemaker. This result can also explain the fact that the lathy shape of the SAN helps to reduce its contact area with the atrium (21). Therefore, in order to achieve the optimal pacemaking capacity, the pacemaker can be moulded into a lathy shape in an isolated (or insulating) way.

### 4.4 Limitations

The present simulation indicated that applying heterotypic gap junctions can improve the drive capacity of bio-pacemaker therapy, but further evidence about the feasibility of constructing heterotypic gap junctions between PM and VM should be verified in biological experiments, and whether the rectified property of gap junctions can improve pacemaker’s drive capacity must be validated in vivo.

Synchrony is a key indicator of ventricular systolic function. The normal ventricle should follow a specific activation sequence. The present simulation was inspired by a judgement about the electronic pacemaker that placing electrodes in the septa was more beneficial for the hemodynamic than in the right ventricular apex (50). In the future, further three-dimensional simulations should be conducted to verify whether the present conclusion can be extended to the whole heart.

## Data Availability Statement

The data that support the findings of this study are available from the corresponding author, [QL and HZ], upon reasonable request.

## Ethical statements

No animal studies are presented in this manuscript.

## Author Contributions

**Conceptualization:** Henggui Zhang.

**Data curation:** Yacong Li.

**Formal analysis:** Yacong Li, Henggui Zhang.

**Funding acquisition:** Qince Li.

**Investigation:** Yacong Li.

**Methodology:** Yacong Li, Jun Liu, Qince Li, Henggui Zhang.

**Project administration:** Lei Ma, Kuanquan Wang, Henggui Zhang.

**Resources:** Lei Ma, Kuanquan Wang, Henggui Zhang.

**Software:** Yacong Li, Jun Liu.

**Supervision:** Lei Ma, Kuanquan Wang, Henggui Zhang.

**Validation:** Yacong Li.

**Visualization:** Yacong Li.

**Writing – original draft:** Yacong Li.

**Writing – review & editing:** Qince Li, Henggui Zhang.

## Supplementary material

Figure S1 Pacemaking cycle length (PCL) and conduction velocity (CV) under different electrical coupling conditions. (A) The PCL of VM tissue under WEC conditions with varying D_PM2VM_ and D_VM2PM_. (B-C) The PCL and CV of VM tissue under W-REC conditions with varying D_PMs_ and D_VM2PM_. White means no spontaneous pacemaking activity.

Figure S2 Potential mapping of the cardiac tissue implanted with a full side-to-side coupling bio-pacemaker distribution. (A) Under original electrical coupling conditions. (B) Under a W-REC condition.

Figure S3 Potential mapping of the cardiac tissue implanted with an embedded bio-pacemaker distribution. (A) Under original electrical coupling conditions. (B) Under a W-REC condition.

Figure S4 Potential mapping of the cardiac tissue implanted with an electrically-isolated bio-pacemaker distribution. (A) In an original electrical coupling condition. (B) In a W-REC condition.

Figure S5 An implantation scheme of a side-to-side coupling bio-pacemaker in idealised cardiac tissue model.

Text S1 Solution of two-dimensional tissue model with nonlinear rectified diffusion coefficient.

Video S1 Spontaneous heartbeats of the isolated bio-pacemaker in real ventricular tissue model.

